# Unveiling Genomic Rearrangements in Engineered iPSC Lines by Optical Genome Mapping

**DOI:** 10.1101/2025.05.10.653237

**Authors:** Darren Finlay, Pooja Hor, Benjamin H. Goldenson, Xiao-Hua Li, Rabi Murad, Dan S. Kaufman, Kristiina Vuori

## Abstract

We demonstrate here the use of optical genome mapping (OGM) to detect genetic alterations arising from gene editing by various technologies in human induced pluripotent stem cells (iPSCs). OGM enables an unbiased and comprehensive analysis of the entire genome, allowing the detection of genomic structural variants (SVs) of all classes with a quantitative variant allele frequency (VAF) sensitivity of 5%. In this pilot study, we conducted a comparative dual analysis between the parental iPSCs and the derived cells that had undergone gene editing using various techniques, including transposons, lentiviral transduction, and CRISPR-Cas9-mediated safe harbor locus insertion at the adeno-associated virus integration site 1 (AAVS1). These analyses demonstrated that iPSCs that had been edited using transposons or lentiviral transduction resulted in a high number of transgene insertions in the genome. In contrast, CRISPR-Cas9 technology resulted in a more precise and limited transgene insertion, with only a single target sequence observed at the intended locus. These studies demonstrate the value of OGM to detect genetic alterations in engineered cell products and suggests that OGM, together with DNA sequencing, could be a valuable tool when evaluating genetically modified iPSCs for research and therapeutic purposes.

## INTRODUCTION

Genetic engineering of cells, including induced pluripotent stem cells (iPSCs), has become a routine practice, with clinical translation of such products advancing steadily. Various techniques are employed to genetically modify cells, including CRISPR-Cas9^1–5^, TALENs^6–9^, and zinc-finger nucleases (ZFNs)^7,10–12^, among others. However, analysis of the engineered cells often focuses primarily on successfully incorporating the desired modification, with limited evaluation of possible other genetic alterations occurring upon genetic engineering. Notably, various genetic modification technologies are known or suspected to cause unintended genomic alterations, such as off-target mutations, genomic rearrangements, or structural variants (SVs)^14,15^. These unintended changes could have significant implications for the safety and efficacy of engineered cell therapies, underscoring the need for a more comprehensive genomic analysis.

In this pilot study, we analyzed the genomic integrity of clinically relevant genetically engineered iPSCs using optical genome mapping (OGM). OGM excels in detecting SVs and providing a comprehensive and unbiased assembly of the entire genome of the sample of interest. Unlike traditional sequencing and computational approaches to genomic SV detection^2,16,17^, OGM offers a distinct advantage due to its capability to analyze significantly longer DNA fragments, typically averaging ∼250 kilobases (kbp) but reaching lengths of up to 2.5 megabases (Mbp). This extended scope allows for a comprehensive and detailed assessment of SVs within the genome, down to a resolution of ∼500 bp. SVs, including insertions, deletions, duplications, and translocations, play a crucial role in understanding genetic diversity, evolution, and disease. The extended fragment length and enhanced resolution of OGM makes it particularly valuable for discerning subtle and large-scale genomic alterations, offering insights into the functional aspects of the genome, and contributing to advancements in various scientific domains, from uncovering disease mechanisms to elucidating evolutionary processes.

iPSCs present both autologous and allogeneic cell therapy avenues for the treatment of various diseases. In most instances, modifying the iPSC genome is essential to customize the cells for particular therapeutic goals. However, iPSCs are known to be susceptible to undesirable genetic modifications not only upon genetic engineering but also during prolonged culture conditions, and there is a notable concern regarding the potential accrual of genomic changes over time^18–21^. Addressing and understanding the dynamics of these genomic SVs is crucial for ensuring the safety and efficacy of iPSC- based cell therapies.

Studies utilizing gene-editing of iPSCs have reported diverse genomic abnormalities, including unexpected copy number losses, chromosomal translocations, and complex genomic rearrangements^3^. Some studies have also documented unexpected large chromosomal deletions (91.2 and 136 kbp) at atypical non-homologous off-target sites, lacking sequence similarity to the single guide RNA (sgRNA) used for CRISPR-Cas9-mediated genome editing^2^. One study that analyzed the impact of prolonged cell culture on iPSCs’ genomic integrity used two related lines that were concurrently subjected to fifty passages. OGM performed at various time points identified several preexisting and culture-acquired SVs, including the acquisition of an extra chromosome 12 in one line^22^. Notably, many of the SVs acquired during culture disrupted protein-coding sequences^2,15^. This underscores the potential pathogenic consequences of undesired SVs induced by genetic engineering, emphasizing the importance of confirming genome integrity before clonal expansion and long-term *ex vivo* culture for both research and clinical applications, as demonstrated in several recent studies^2,22–26^. Thus, balancing the benefits of allogeneic cell therapies with the potential risks associated with genetic modifications highlights the importance of rigorous genomic monitoring and validation throughout the iPSC culture and engineering processes to develop safe and effective cell therapies.

We evaluate here the use of OGM to quantify and monitor genetic alterations in iPSCs that have been engineered to derive immune cells with improved anti-tumor activity. We conducted a dual analysis to compare genetically edited progeny modified through transposons, lentivirus, or CRISPR-Cas9-mediated safe harbor locus insertion at the AAVS1 site with their corresponding wildtype (WT) parental iPSC lines. These parental lines had been derived from various sources, including CD34+ cells from umbilical cord blood (UCB) and peripheral blood mononuclear cells (PBMCs). Our studies demonstrate that editing iPSCs with lentiviral transduction or transposons resulted in an elevated number of transgene insertions in each clonal line, exceeding the desired quantity of insertions. In contrast, CRISPR-Cas9-mediated safe harbor locus insertion exhibited highly precise outcomes, with only a single insertion observed in the intended locus. Furthermore, lentiviral or transposon-mediated editing revealed many unique genomic SVs in the edited iPSCs that were absent in the parental line. Comparative analysis with the safe harbor insertion line, in turn, showed no undesired SVs in one sample and only one other unique SV (a duplication) in another.

In sum, we demonstrate that OGM can provide a genome-wide understanding of genomic insertions and rearrangements occurring upon gene editing, and in a manner that is faster, more economical, and more precise than conventional cytogenetics methods. Our intent here is not to compare and contrast various genetic engineering technologies for their utility. Indeed, robust functional studies beyond the scope of this work are required for careful selection of editing technologies and edited cells to ensure their safety and efficacy in research and therapeutic applications. Rather, we propose that OGM, when combined with DNA sequencing, can serve as a useful tool in the development and optimization of cell therapy products, offering valuable insights into the genetic integrity of edited cells and supporting their selection for safe and effective therapeutic use.

## RESULTS

### Description of the Optical Genome Mapping process

To investigate potential genomic rearrangements in our engineered iPSC samples, we employed optical genome mapping (OGM), as illustrated in Figure 1, to identify genomic structural variants (SVs). The Bionano platform (Saphyr) was utilized for OGM analysis, allowing for the assessment of genomic integrity in both parental and engineered iPSC lines. The experimental process involved several sequential steps starting with isolating high molecular weight DNA from target cells. Subsequently, sequences across the entire genome were fluorescently labeled, followed by the transfer of the labeled DNA into a microchip for analysis (see Materials and Methods). Through repeated cycling of labeled, linearized DNA, the labeled genome was scanned. Utilizing Bionano Access (v1.8.1), consensus optical genome maps were assembled, facilitating easy visualization of genome alterations and enabling the identification of chromosomal aberrations and structural variants (Fig. 1). Figure 1B depicts a generic Circos plot output labelled to describe each of the output parameters.

**Figure 1.**
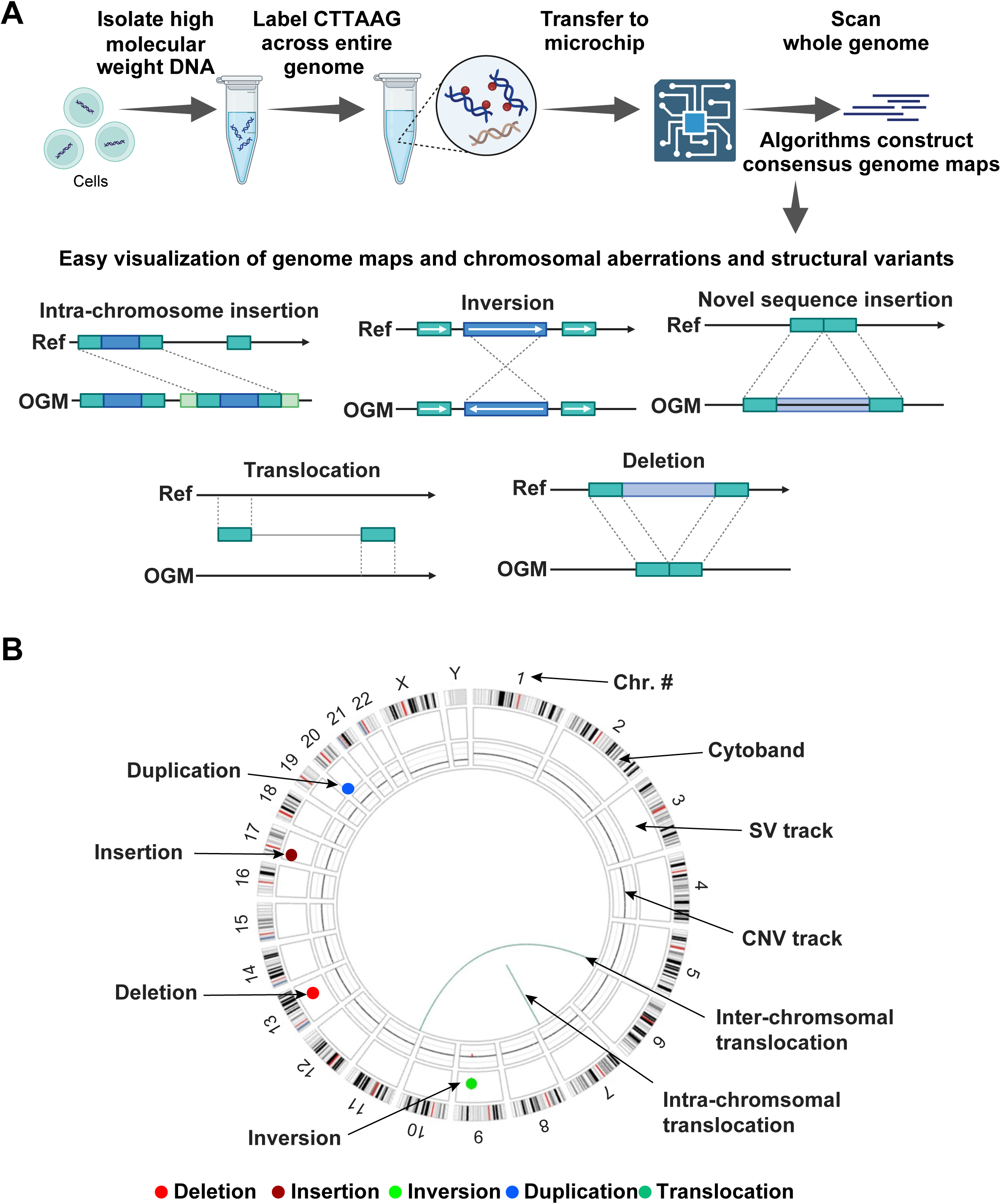
(A) Schematic of the experimental flowchart for utilizing optical genome mapping to detect genomic SVs and to evaluate the genomic integrity of genetically modified iPSC lines in comparison to the parental iPSC lines. (B) Generic Circos plot depicting what each ring, and colored SV “spot”, represents. The outer ring shows cytobands and chromosome number (Chr. #). The next inner ring shows the SV track where different colored spots depict distinct SVs. The next inner ring depicts the copy number variation (CNV) track, which is normal (and one copy each of X and Y chromosomes) for this schematic, while the central lines (teal) represent intra- and inter-chromosomal translocations. Color scheme of detected SVs is consistent throughout the manuscript, bright red = deletion; burgundy = insertion; blue = duplication; green = inversion; and teal = intra-fusions/ translocations.

In order to fully understand the true genome status of the parental cell models utilized in this study, we first analyzed them using the Rare Variant Analysis (RVA) pipeline. As the name suggests, the RVA pipeline is capable of detecting SVs with a quantifiable detection rate of SVs at an allele frequency of 5%. We note, however, that in our experience, the RVA pipeline is sometimes capable of detecting SVs of less than 1% allele frequency, although non-quantifiably. These outputs also quantify the variant allele frequency (VAF), which can be considered a measure of clonality. Furthermore, the detected SVs are filtered to remove any variant that has ever been detected in a cohort of healthy volunteers, and we have further filtered the dataset to describe only SVs that occur within 12kbp of a canonical protein-coding gene as annotated in the latest human genome build (GRCh38/ hg38).

As described below, we then examined the effectiveness of OGM analyses to evaluate distinct structural variations by comparing parental iPSC lines with their genetically modified iPSC counterparts that had been edited using widely used genome editing tools: CRISPR-Cas9, transposons (piggyBac), and lentivirus.

### Dual analysis comparing parental iPSCs and their genetically modified progeny employing CRISPR-Cas9 insertion at the AAVS1 safe harbor site

Dual analysis feature of Bionano Solve (v3.8.1) facilitates a direct comparison between the genetically modified cell line and its parental counterpart. Such an analysis filters out any SVs common to both samples to reveal *<u>only those SVs unique to the engineered</u> <u>cells</u>*. Figure 2 demonstrates this dual analysis comparison between CRISPR-Cas9 edited iPSCs with their parental cells. Using the CRISPR-Cas9 method, we genetically modified two individual WT iPSCs, inserting two different chimeric antigen receptor (CAR) constructs (anti-Mesothelin CAR and anti-CD123-IL15 CAR) into the AAVS1 safe harbor locus. The parental WT iPSC samples used in this study had been generated via episomal reprogramming using Sendai virus^27^ from umbilical cord blood (UCB) or peripheral blood mononuclear cells (PB). The anti-mesothelin (Meso) CAR construct was engineered into the WT PB-iPSC parental line, resulting in AAVS1-Meso-Bai1 line. Whereas, the anti-

**Figure 2.**
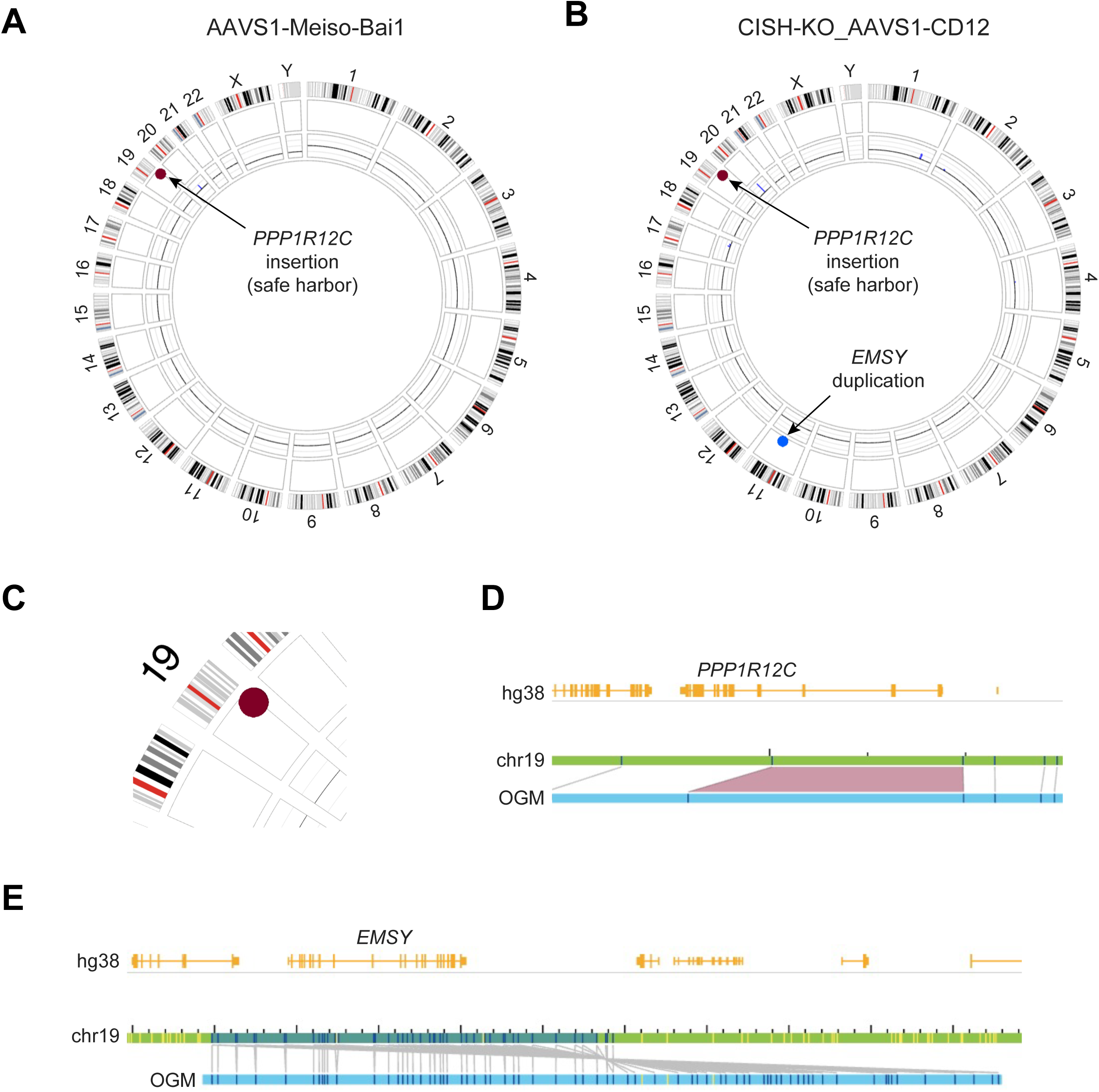
Dual analysis comparing iPSCs engineered by CRISPR-Cas9 for safe harbor locus insertion at the AAVS1 site with their parental counterparts. (A) Circos plot, showing *only SVs unique* to the genetically modified cells, of AAVS1-Meso-Bai1 iPSC (vs. WT PB- iPSC). (B) Circos plot depicting *only unique SVs* of CISH-KO AAVS1-CD123-IL15 iPSC (vs. CISH-KO iPSC). (C) Zoomed in view of AAVS1-Meso-Bai1 iPSC Chr. 19 insertion (ins(19;?)(q13.42;?)). (D) Gene level map showing insertion in the safe harbor locus in *PPP1R12C* of AAVS11-Meso-Bai1 iPSC in Chr. 19. E) Gene level map showing inverted duplication of *EMSY* gene. Gene level maps show (from top to bottom): the annotated gene(s) (hg38) in yellow, the “parental” iPSC chromosome(s) (chr#) in green, and the optical genome map (OGM) in blue. Dark blue ticks represent detected labels, and the light grey lines represent matching labels. The purple trapezoid(s) represents an insertion or deletion, where the labels are further, or closer, apart respectively than in the parental sample.

CD123-IL15 CAR construct was engineered into WT UCB-iPSC line, which additionally had been previously engineered to knockout Cytokine-inducible SH2-containing protein (CIS; encoded by the gene *CISH*)^28^, resulting in CISH-KO AAVS1-CD123-IL15 line. Figure 2A shows a Circos plot of only unique SVs in AAVS1-Meso-Bai1 iPSCs when compared against their parental (WT PB-iPSC) cells. The only SV observed is an insertion in Chr. 19 within the expected safe harbor locus ((q13.42) in *PPP1R12C*) (Fig. 2C and D). Safe harbor insertion is also detected in CISH-KO AAVS1-CD123-IL15 iPSCs that is not present in the parental (CISH-KO iPSC) cells (Fig. 2B), but these cells also contain a 234.5kbp duplication in Chr. 11. This inverted duplication (Fig. 2E) results in a duplication of the entire *EMSY* gene, encoding EMSY, a protein known to interact with BRCA2^29,30^.

### Dual analysis comparing parental iPSCs with their genetically modified progeny utilizing the piggyBac transposon system

We performed similar analyses on three genetically modified iPSC samples that had been engineered using the piggyBac transposon system. The anti-CLL1 CAR-IL15 receptor fusion (anti-CLL1-IL15RF), switchable-CAR (sCAR), and anti-Mesothelin CAR (anti- Meso) constructs were cloned into the piggyBac transposon vector. The first two CAR constructs were engineered into WT UCB-iPSC line and the anti-Meso CAR was engineered into WT PB-iPSC line. Figure 3 depicts Circos plots illustrating the unique SVs alongside graphs plotting SV size against VAF for: A) anti-CLL1-IL15RF iPSC, B) anti-Meso iPSC, and C) sCAR iPSC in comparison to their respective parental WT iPSC lines. A significantly higher number of SVs were detected in these three engineered cells generated utilizing the piggyBac system when compared to those observed with CRISPR- Cas9 safe harbor (AAVS1) locus insertion (Fig. 2). We detected 46 insertions in the anti- CLL1-IL15RF iPSCs, 8 in the anti-Meso iPSCs, and 19 in the sCAR iPSCs engineered by the piggyBac system. Intriguingly, most of the insertions in each sample were of very similar sizes (∼6.8kbp for anti CLL1-IL15RF iPSC, ∼12.2kbp for anti-Meso iPSC, and ∼10.9kbp for sCAR iPSC; Fig. 3A-C, lower panels, respectively) that correspond with the sizes of the CAR constructs used, suggesting that they are the desired insertion sequence in each case. Furthermore, all these piggyBac insertion events are dispersed throughout the genome. As such, multiple genes per sample could in principle be perturbed by these insertions. Interestingly, the similarly sized insertions have a wide variation in VAF as seen in the presented SV size versus VAF graphs. While several of the similarly sized insertions have VAFs of ∼0.5, many have a much lower frequency of insertion (e.g., Fig. 3B), suggesting widespread clonality in the cell population in question. In addition to the similar sized inversions, all three samples also possess larger insertion SVs, with some deletions, a duplication, and a translocation. In fact, the anti-CLL1-IL15RF iPSCs’ translocation is predicted to result in a putative gene fusion between the genes *AHI1* and *SYNE1* (fus(6;6) (q23.3;q25.2)) (Fig. 3D).

**Figure 3.**
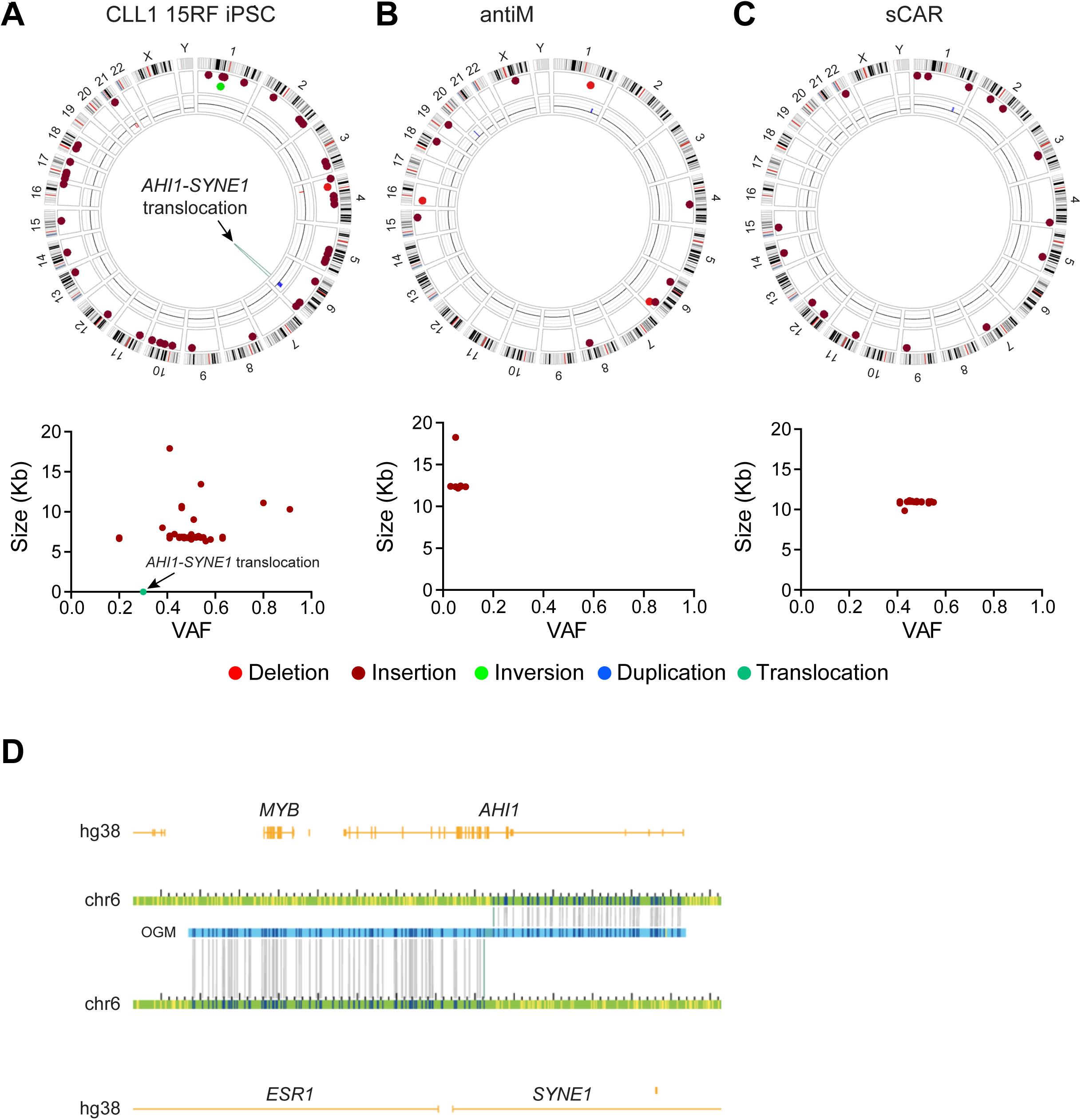
Dual analysis comparing piggyBac transposon edited iPSCs with their parental counterparts. (A) Upper panel, Circos plot of only unique SVs of CLL1-IL15RF iPSC (vs. WT UCB-iPSC); lower panel, plot of VAF vs. SV size. (B) Upper panel, Circos plot of only unique SVs of anti-Meso CAR iPSC (vs. WT PB-iPSC); lower panel, plot of VAF vs. SV size. (C) Upper panel, Circos plot of only unique SVs of sCAR iPSC (vs. WT UCB-iPSC); lower panel, plot of VAF vs. SV size. D) Gene level map of the predicted gene fusion of *AHI1* and *SYNE1* (fus(6;6)(q23.3;q25.2)).

### Dual analysis comparing parental iPSCs with their genetically modified progeny utilizing lentiviral system

To further investigate potential genomic abnormalities in engineered iPSCs we also performed OGM and dual analysis on samples that had been genetically edited using a lentiviral system. Two different CAR constructs, anti-CLL1 and anti-CD123, were cloned into the pLenti vector backbone. Parental WT UCB-iPSCs and PB-iPSCs were transduced respectively with the lentivirus encoding for anti-CLL1-CAR and anti-CD123- CAR and clonally propagated. Figure 4 illustrates Circos plots, and graphs depicting SV size versus VAF, after dual analysis of lentiviral introduction for A) anti-CD123-CAR iPSCs, and B) anti-CLL1-CAR iPSCs, compared to their respective parental cells. The anti-CD123-CAR iPSCs have 105 high-confidence gene-overlapping insertions. 103 of those insertions are ∼7kbp (7073.46 +/- 235.46bp), and the other two are ∼14kbp, and as with the non-clonally selected transposon system sample (Fig. 4B), there is a wide spread of associated VAF (lower panel) with most detected insertions at less than 0.25. Furthermore, this lentivirally modified iPSC population also has two large, deleted regions; a 225.7kbp region on Chr. 15 and a 121.4kbp region on Chr. 17 (Fig. 4A).

**Figure 4.**
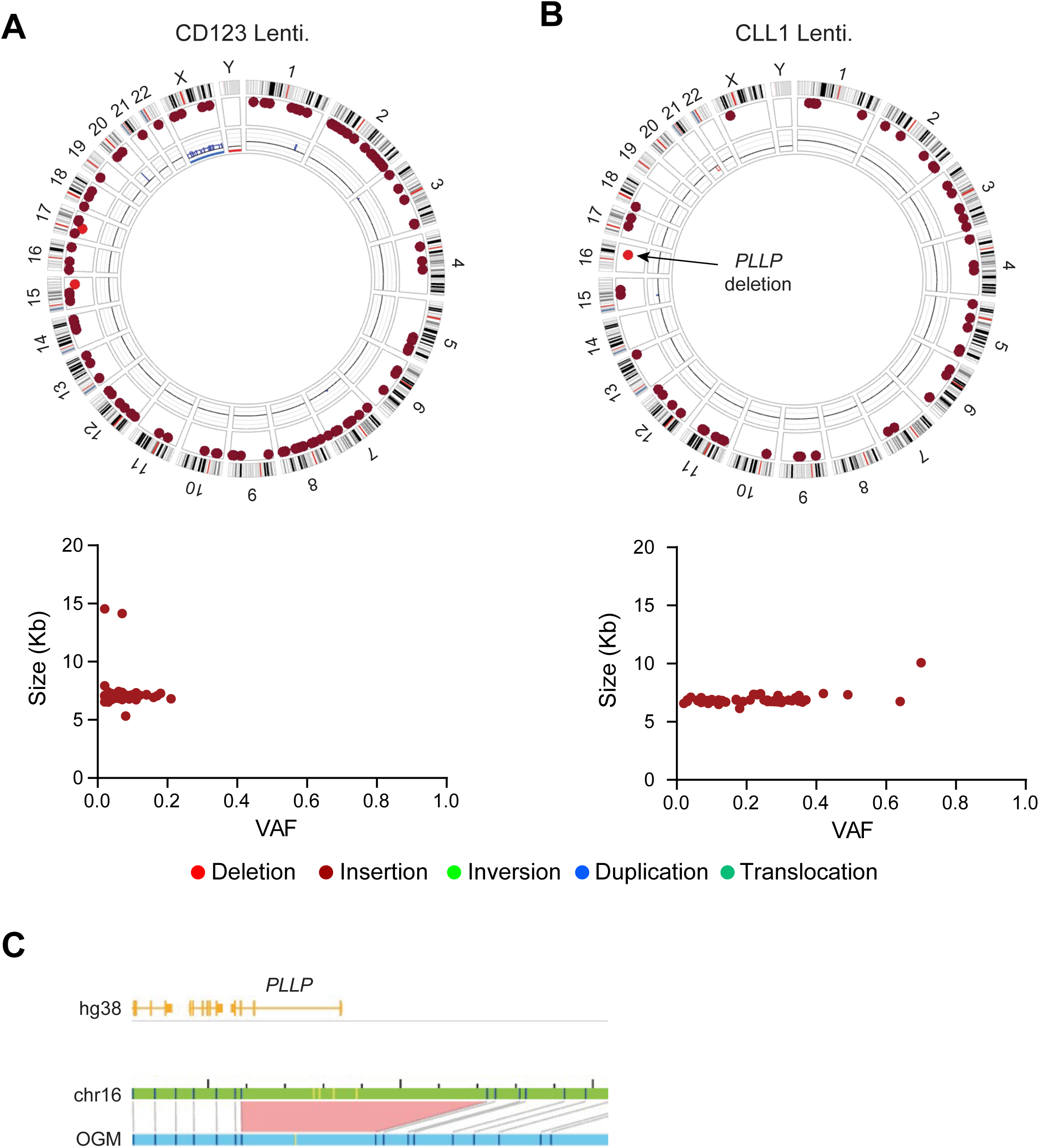
Dual analysis comparing lentivirally edited iPSCs with their parental cells. Upper panels show Circos plots of SVs and lower panels show plots of VAF vs. SV size of (A) CD123-CAR iPSC and (B) CLL1-CAR iPSC (both vs. parental WT UCB-iPSCs). (C) Gene level map of *PLLP* deletion event.

The dual analysis of the second lentivirally engineered cell sample, anti-CLL1-CAR iPSCs (Fig. 4B), demonstrates that these cells possess 51 insertions not present in the parental cells, again spread throughout the genome, and with differing VAFs (Fig. 4B, lower panel). All of these insertions are ∼7kbp (6867.31 +/- 283.40bp, n=51) except for one 10067bp insertion in the gene *ADGRF3*, encoding Adhesion G Protein-Coupled Receptor F3. These cells also have a deletion (29.1kbp) within the *PLLP* gene, although the VAF for this SV is only 0.12 (Fig. 4C). Details of all unique dual analysis detected SVs are provided as Supplemental Data S1.

### Evaluating OGM analysis proficiency in detecting gene knockouts in genetically modified iPSCs

The OGM platform demonstrates proficiency in identifying large or challenging-to-detect SVs^31–35^. Here, we conducted further analysis to assess its capability in detecting smaller deletions within the genome. Genetically modified iPSC samples harboring intended small gene knock-out deletions were therefore subjected to OGM analysis. Figure 5 illustrates the dual analysis outputs from (A) HIF1A-KO iPSCs and (B) CISH-KO iPSCs^28^, compared to their respective parental cells. Both samples underwent knockout confirmation through PCR and DNA sequencing. The confirmation data for CISH-KO iPSCs has been previously published^24^. Notably, the OGM analysis is not able to detect the desired 60bp deletion in the *HIF1A* gene on Chr. 14 (Fig. 5A), while it was confirmed using PCR as expected. Similarly, no indels were detected in the *CISH* gene (exon 3, 3067-3185 bp region) on Chr. 3 (Fig. 5B), while such small indels were confirmed by Sanger sequencing in previously published work^28^. However, the HIF1A-KO iPSC cells did exhibit two apparently heterozygous deletions not observed in the parental cells (250.6kbp in Chr. 1 and 53.4kbp in Chr. 16, Fig. 5A). The Chr. 16 deletion results in an SV within *EEF2K*, encoding Eukaryotic Elongation Factor 2 Kinase (Fig. 5C).

**Figure 5.**
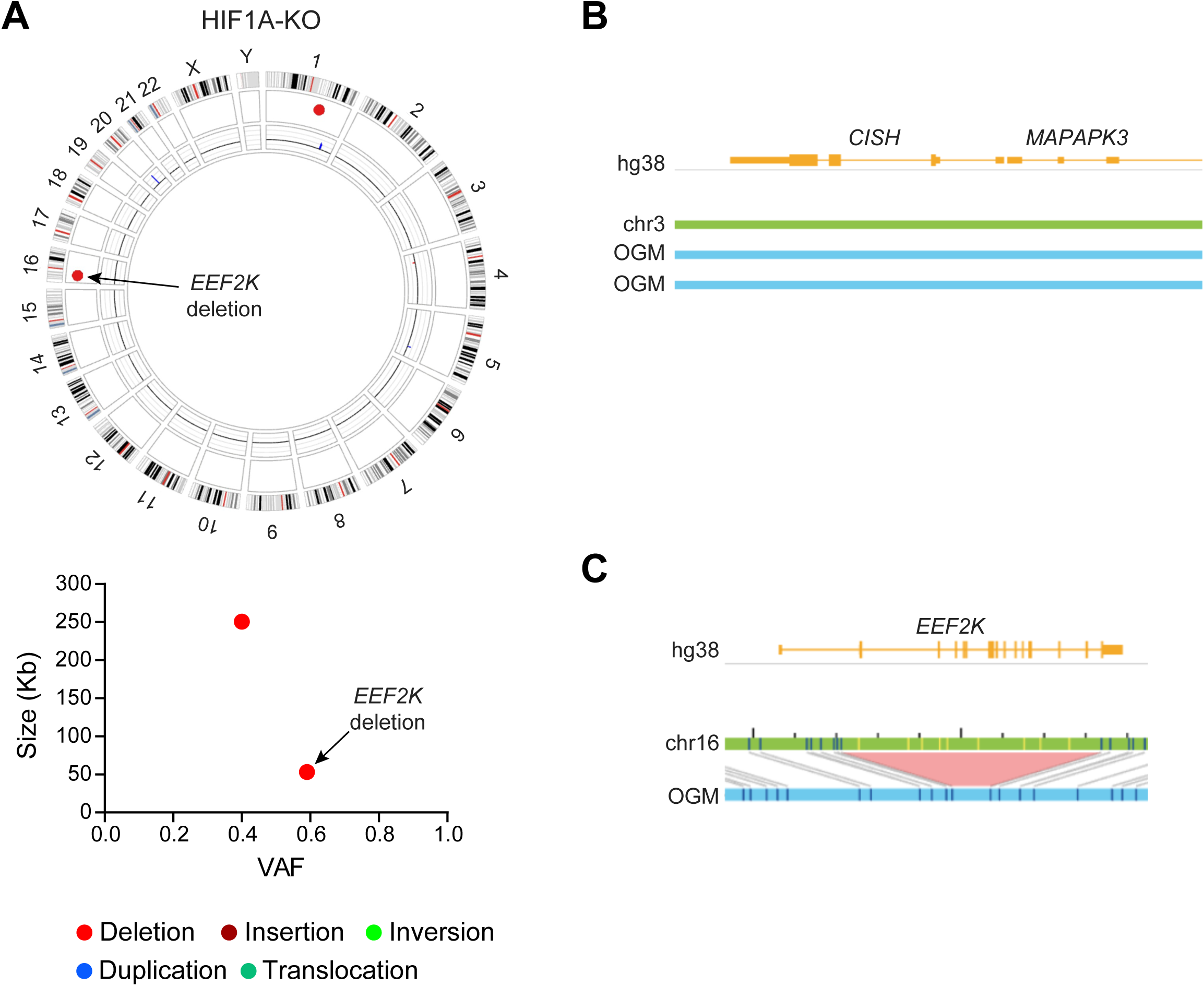
Dual analysis comparison is incapable of detecting gene knockout events smaller than ∼500 base pairs. (A) Upper panel, Circos plot of HIF1A-KO iPSC unique SVs (vs. parental WT UCB-iPSC) shows that OGM fails to detect the ∼60bp HIF1A deletion in Chr. 14, and lower panel, plot of VAF vs. SV size. (B) Gene level map of *CISH* gene locus in CISH-KO iPSC shows no detected deletion. (C) Gene level map of a deletion event in the *EEF2K* gene in HIF1A-KO iPSC cells.

In conclusion, our study demonstrates OGM’s efficacy in identifying SVs in parental cells and detecting both desired and undesired SVs in their engineered variants. Our findings indicate that CRISPR-Cas9 safe harbor (AAVS1) site insertion yields only the intended insertion, whereas transposon or lentiviral techniques lead to a higher frequency of insertions and off-target genomic SV events, potentially of biological and clinical significance. For detection of SVs less than approximately 500 bp, complementary DNA sequencing would be required.

## DISCUSSION

Gene editing tools have revolutionized cell and gene therapy by enabling precise modifications to the genome, offering new avenues for treating genetic disorders and diseases. These tools, such as CRISPR-Cas9, transposons, and lentiviral delivery systems, hold promise in the development of personalized therapies, enhancing the efficiency and safety of gene therapy approaches.

Viral delivery systems, such as retrovirus and lentivirus, are now commonly used for clinical cell engineering. They integrate into the host genome, providing stable and long-term transgene expression of the therapeutic gene. Additionally, they can carry cargo anywhere up to 8-10kbps for lentivirus and 5-7kbps for retrovirus, making them suitable for a variety of therapies^36–38^. Beyond viral systems, non-viral methods such as PiggyBac transposon and CRISPR-Cas9 genome editing are gaining traction due to their versatility and reduced risk of insertional mutagenesis. PiggyBac enables stable transgene integration without viral vectors and is particularly advantageous for applications requiring seamless transgene excision^39,40^. Meanwhile, CRISPR-Cas9 has revolutionized gene editing by providing precise modifications at target loci, though challenges such as off- target effects and delivery efficiency remain areas of active research^41,42^. Emerging innovations, including base editing and prime editing, aim to refine genome modifications with higher precision and fewer unintended mutations, further expanding the toolbox for genetic engineering^43^.

Genetically modified iPSC-derived cells offer potential as allogeneic resources for cell therapy, providing a scalable and standardized approach to treatment. Different genomic analysis methods are utilized to detect chromosomal abnormalities, each with its own advantages and limitations. Karyotyping, a traditional cytogenetic technique, provides a genome-wide view but suffers from very low resolution, high subjectivity, and an inability to detect allelic imbalances. Additionally, it requires a complex workflow, specialized personnel, and cell culturing, leading to extended turnaround times of ∼1-2 weeks in-house and up to ∼5-8 weeks when outsourced^44^. Polymerase chain reaction (PCR) based methods offer a faster alternative, typically completed in under a week; however, they are limited to detecting aneuploidies and to specific target sites rather than providing a genome-wide analysis^45^. Chromosomal microarray analysis (CMA) improves on resolution and it allows for the detection of copy number variations (CNVs) and allelic imbalances, but it is unable to identify balanced rearrangements and struggles with repeat-rich regions^46^. Next-generation sequencing (NGS) offers a powerful platform for high-throughput genomic analysis, but its effectiveness varies depending on the approach. Targeted sequencing does not reliably detect CNVs or structural variants (SVs), and whole-genome sequencing, while comprehensive, is costly and has low sensitivity for CNVs and SVs in heterogeneous samples. Furthermore, the time required for NGS-based analysis is variable, depending on the specific sequencing depth and bioinformatics pipelines^47^.

Based on these observations we decided to study the utility of OGM to assess genomic integrity and the potential acquisition of undesired SVs in various genetically modified iPSC model systems. Our studies provide further support for the utility of OGM in offering a rapid, cost-effective, unbiased, and efficient means of screening for genes affected by structural variations (SVs) in the context of genetically modified iPSCs. OGM assembles an unbiased, genome-wide ’optical map’ of a sample of interest which can be compared to a reference sequence of choice to assess alterations in genome integrity (Fig. 1). When comparing the assembled genome from an iPSC model to the Human Genome Project’s reference build (GRCh38/hg38), as in Rare Variant Analysis (RVA), it can detect SVs at a resolution of 500 bp with a quantitative variant allele fraction (VAF) of 5%, thus allowing detection of rare SVs and providing an output of VAF, a measure of clonality of the event. In this analysis, any SV ever detected in a database of healthy volunteers is removed to only show these rare variants. We note that in this study, we further filtered all SVs to present and discuss only those events that are associated with canonical protein-coding genes as defined by GRCh38/hg38. Therefore, the total number of insertions and off-target genomic perturbations is, in fact, far greater than has been presented herein.

To focus exclusively on SVs only present in the genetically modified cells and not in the parental lines, we utilized RVA dual analysis to filter out shared SVs and used these outputs to evaluate genomic alterations resulting from three different genetic modification methods. Our intent here is not to compare and contrast various genetic engineering methods for their robustness or utility. Rather, our goal is to provide proof-of-concept that OGM can provide a digital workflow, allowing for efficient and reproducible analysis of large genome datasets, facilitating faster quality control processes for edited cells. Nevertheless, some general observations can be made. CRISPR-Cas9 targeted insertion in the AAVS1 safe harbor site resulted in an insertion only at that locus in one iPSC sample (Fig. 2A, B, and C). Another sample also had a single insertion at the same locus on Chr. 19, however, this sample additionally had a duplication on Chr. 11 (Fig. 2B and Suppl. Data S1). Of note, it is unclear if this duplication occurred due to the CRISPR-Cas9 editing or if it was acquired during subsequent selection and propagation stages of the cells. The use of transposons or lentivirus methodologies resulted in multiple insertions of genetic material throughout the iPSC genome (Figs. 3 and 4). For example, a sample modified using lentivirus (anti-CD123 iPSC, Fig. 4A) has 105 unique insertions, almost all of which (103/105) are ∼7kbp suggesting this is the desired insertion cassette (Fig. 4A, lower panel). Similarly using the piggyBac transposon system we also detected multiple insertions throughout the genome within canonical genes (Fig. 3). As an example, one piggyBac-modified sample (CLL1-IL15RF iPSC, Fig. 3A) has 46 insertions. It also has a deletion, an inversion, and an intra-chromosomal translocation event. Indeed this Chr. 6 translocation event is predicted to result in a novel fusion of the genes *AHI1* and *SYNE1* (Fig. 3D). *AHI1* acts as a sensor for insulin signaling and is vital for both cerebellar and cortical development in humans. *SYNE1* encodes the Syne-1 protein, which plays a role in maintaining the cerebellum, the brain region responsible for movement control. Variants in both genes have been linked to various neurological and neuromuscular disorders, suggesting important cellular functions for these gene products. Consequently, a fusion of these two genes may lead to unforeseen ramifications, potentially disrupting their normal functions and/ or resulting in aberrant cellular events.

In summary, we demonstrate the use of OGM to detect genomic structural variants (SVs) in human iPSCs in an unbiased, genome-wide manner. Through dual analyses, we not only identify the desired insertion sequences in engineered iPSC lines but also uncover unintended off-target SVs. While the exact effects of both desired and off-target genomic changes on cellular function remain a focus of our future research, our findings highlight the need for rigorous assessment of genetic fidelity in both parental and genetically modified iPSC lines, particularly as these cells transition to clinical use. Obviously, many other bottlenecks to clinical application exist, such as the high costs involved with clinical cell production, limited cargo capacity, and appropriate regulatory requirements. Within the scope of the current study, we propose that OGM offers a cost- effective, scalable, and streamlined approach for detecting genomic integrity compared to other analysis methods, making it a potentially invaluable tool for clinical products that combine cell and gene engineering workflows.

## MATERIALS AND METHODS

### iPSC cells and culture

hiPSCs were derived from CD34+ cells obtained from peripheral blood (PB) or umbilical cord blood (UCB), sourced from different donors, and maintained in a feeder-free manner following established protocols^48–50^. hiPSCs were passaged with Accutase (STEMCELL Technologies, Vancouver, Canada, Cat. 07920) at a 1:4 to 1:10 ratio on Matrigel-coated plates (Corning, NY, U.S., Cat. no. 354277), using mTeSR™ Plus media (STEMCELL Technologies, Vancouver, Canada, Cat. no. 100-0276). Cells were passaged when they reached approximately 85-90% confluency.

### CRISPR-Cas9 AAVS1 safe harbor insertions

The Bai1-CAR-Zeo::EGFP and CD123-CAR-IL15-Zeo::EGFP constructs were cloned into the AAVS1-Pur-CAG-EGFP donor plasmid under the CAG promoter^51–53^. RNP complex constituting the donor plasmid, Cas9 protein (IDT), and gRNA (GGGGCCACTAGGGACAGGAT) was nucleofected into the wild-type iPSCs using the Amaxa® 4D-Nucleofector® Basic Protocol for Human Stem Cells. The positive clones were selected through puromycin and/or zeocin treatment because the engineered iPSC lines conferred dual drug resistance. This was due to the inserts encoding a zeocin (bleomycin) resistance cassette and the donor plasmid expressing a Puromycin resistance cassette.

### CRISPR - Cas9 gene knockout generation

The detailed method for generating the CISH-KO iPSC line is documented in our previously published work^28^. The HIF1A-KO iPSC line was generated using CRISPR- Cas9 technology with a pair of guide RNAs targeting exon 1 of the HIF-1a gene in the UCB-iPSCs.

### piggyBac transposon

The wild-type iPSCs underwent re-engineering through the introduction of a piggyBac transposon vector expressing anti-CLL1-CAR-IL15, anti-Mesothelin-CAR, or anti- PNEscFV-CAR4 NK-CAR constructs, resulting in the generation of transgenic iPSC lines^51,54,55^.

### Lentiviral systems

The transgenic iPSC lines were generated by introducing lentiviral insertions using molecular constructs anti-CD123-CAR and anti-CLL1-CAR in the pLenti vector backbone^52,53,56^.

### Optical Genome Mapping (OGM)

OGM was carried out using the Saphyr system (Bionano Genomics Inc., San Diego, CA) essentially as described previously^33^. Briefly, ultra-high molecular weight (UHMW) DNA was isolated from 1- 1.5M cells of interest using the Bionano Prep SP Blood and Cell Culture DNA Isolation Kit (#80030) exactly as per the manufacturer’s instructions. 750 ng of purified and homogeneous UHMW DNA was fluorescently labeled genome wide on a 6bp motif (CTTAAG, ∼14-17 labels per 100kbp of human genome) using Bionano’s Direct Label and Stain (DLS) kit (#80005), exactly as per the manufacturer. Purified labelled DNA was loaded onto G2.3 Saphyr Chips (#20366) and run on the Saphyr until >1300 Gbp worth of DNA was mapped (>75% map rate = >300x coverage).

### Rare Variant Analysis (RVA) and Dual Analysis

To call rare, or unique, SVs the rare variant analysis (RVA) pipeline and the dual analysis features, respectively, of the Bionano software were used. Specifically, we used Bionano Access v1.8.1 and Bionano Solve v3.8.1 and GRCh38 (Genome Reference Consortium Human Build 38) as reference sequence for RVA. SVs were only called if a minimum of 5 DNA molecules spanning each SV breakpoint were detected and assessed. Only SVs with a confidence call of >0.99 were considered. RVA detected SVs were further filtered to remove any variants ever observed in a control database of 179 healthy individuals. For the analyses presented herein, only SVs that occurred within 12kbp of an annotated canonical protein-coding gene are shown.

## DATA AVAILABILITY

All Dual Analysis detected (canonical gene associated) SVs can be found in Supplemental Data S1. Original/ raw data files are available on request (contact).

## ACKNOWLEDGEMENTS

The authors thank SBP’s Tumor Analysis, and Bioinformatics, Shared Resources for OGM analyses and Figure preparation. All SBP Shared Resources are supported by SBP’s NCI Cancer Center Support Grant, P30 CA030199. UCSD studies were supported by the NIH/NCI grants U01CA217885, P30CA023100 (administrative supplement) and the Sanford Stem Cell Institute at UCSD.

## AUTHOR CONTRIBUTIONS

Darren Finlay: Conception and design, collection and/or assembly of data, data analysis and interpretation, manuscript writing.

Pooja Hor: Conception and design, provision of study material, manuscript writing.

Benjamin H. Goldenson: Provision of study material.

Xiao-Hua Li: Provision of study material.

Rabi Murad: Collection and/or assembly of data, other (figure preparation).

Dan S. Kaufman: Conception and design, financial support, data analysis and interpretation, manuscript writing, final approval of manuscript.

Kristiina Vuori: Conception and design, financial support, data analysis and interpretation, manuscript writing, final approval of manuscript.

---

Supplemental Data S1. Excel file of Dual Analysis of each unique SV not present in parental cells detected. Only SVs within 12kbp of a canonical gene are presented for clarity.

## REFERENCES

1. Ran, F.A., Hsu, P.D., Wright, J., Agarwala, V., Scott, D.A., and Zhang, F. (2013). Genome engineering using the CRISPR-Cas9 system. Nat Protoc 8, 2281–2308. 10.1038/nprot.2013.143.

2. Tsai, H.-H., Kao, H.-J., Kuo, M.-W., Lin, C.-H., Chang, C.-M., Chen, Y.-Y., Chen, H.-H., Kwok, P.-Y., Yu, A.L., and Yu, J. (2023). Whole genomic analysis reveals atypical non-homologous off-target large structural variants induced by CRISPR- Cas9-mediated genome editing. Nat Commun 14, 5183. 10.1038/s41467-023-40901-x.

3. Kitano, Y., Nishimura, S., Kato, T.M., Ueda, A., Takigawa, K., Umekage, M., Nomura, M., Kawakami, A., Ogawa, H., Xu, H., et al. (2022). Generation of hypoimmunogenic induced pluripotent stem cells by CRISPR-Cas9 system and detailed evaluation for clinical application. Molecular Therapy Methods & Clinical Development 26, 15–25. 10.1016/j.omtm.2022.05.010.

4. Giacalone, J.C., Sharma, T.P., Burnight, E.R., Fingert, J.F., Mullins, R.F., Stone, E.M., and Tucker, B.A. (2018). CRISPR-Cas9 Based Genome Editing of Human Induced Pluripotent Stem Cells. Curr Protoc Stem Cell Biol 44, 5B.7.1–5B.7.22. 10.1002/cpsc.46.

5. Kosicki, M., Tomberg, K., and Bradley, A. (2018). Repair of double-strand breaks induced by CRISPR-Cas9 leads to large deletions and complex rearrangements. Nat Biotechnol 36, 765–771. 10.1038/nbt.4192.

6. Cermak, T., Doyle, E.L., Christian, M., Wang, L., Zhang, Y., Schmidt, C., Baller, J.A., Somia, N.V., Bogdanove, A.J., and Voytas, D.F. (2011). Efficient design and assembly of custom TALEN and other TAL effector-based constructs for DNA targeting. Nucleic Acids Res 39, e82. 10.1093/nar/gkr218.

7. Gaj, T., Gersbach, C.A., and Barbas, C.F. (2013). ZFN, TALEN, and CRISPR/Cas-based methods for genome engineering. Trends Biotechnol 31, 397–405. 10.1016/j.tibtech.2013.04.004.

8. Ding, Q., Lee, Y.-K., Schaefer, E.A.K., Peters, D.T., Veres, A., Kim, K., Kuperwasser, N., Motola, D.L., Meissner, T.B., Hendriks, W.T., et al. (2013). A TALEN Genome-Editing System for Generating Human Stem Cell-Based Disease Models. Cell Stem Cell 12, 238–251. 10.1016/j.stem.2012.11.011.

9. Smith, C., Gore, A., Yan, W., Abalde-Atristain, L., Li, Z., He, C., Wang, Y., Brodsky, R.A., Zhang, K., Cheng, L., et al. (2014). Whole-Genome Sequencing Analysis Reveals High Specificity of CRISPR/Cas9 and TALEN-Based Genome Editing in Human iPSCs. Cell Stem Cell 15, 12–13. 10.1016/j.stem.2014.06.011.

10. Urnov, F.D., Rebar, E.J., Holmes, M.C., Zhang, H.S., and Gregory, P.D. (2010). Genome editing with engineered zinc finger nucleases. Nat Rev Genet 11, 636–646. 10.1038/nrg2842.

11. Hockemeyer, D., Soldner, F., Beard, C., Gao, Q., Mitalipova, M., DeKelver, R.C., Katibah, G.E., Amora, R., Boydston, E.A., Zeitler, B., et al. (2009). Efficient targeting of expressed and silent genes in human ESCs and iPSCs using zinc-finger nucleases. Nat Biotechnol 27, 851–857. 10.1038/nbt.1562.

12. Soldner, F., Laganière, J., Cheng, A.W., Hockemeyer, D., Gao, Q., Alagappan, R., Khurana, V., Golbe, L.I., Myers, R.H., Lindquist, S., et al. (2011). Generation of Isogenic Pluripotent Stem Cells Differing Exclusively at Two Early Onset Parkinson Point Mutations. Cell 146, 318–331. 10.1016/j.cell.2011.06.019.

13. Gaj, T., Sirk, S.J., Shui, S., and Liu, J. (2016). Genome-Editing Technologies: Principles and Applications. Cold Spring Harb Perspect Biol 8, a023754. 10.1101/cshperspect.a023754.

14. Zuccaro, M.V., Xu, J., Mitchell, C., Marin, D., Zimmerman, R., Rana, B., Weinstein, E., King, R.T., Palmerola, K.L., Smith, M.E., et al. (2020). Allele-Specific Chromosome Removal after Cas9 Cleavage in Human Embryos. Cell 183, 1650–1664.e15. 10.1016/j.cell.2020.10.025.

15. Kosicki, M., Tomberg, K., and Bradley, A. (2018). Repair of double-strand breaks induced by CRISPR-Cas9 leads to large deletions and complex rearrangements. Nat Biotechnol 36, 765–771. 10.1038/nbt.4192.

16. Demidov, G., Laurie, S., Torella, A., Piluso, G., Scala, M., Morleo, M., Nigro, V., Graessner, H., Banka, S., Lohmann, K., et al. (2024). Structural variant calling and clinical interpretation in 6224 unsolved rare disease exomes. Eur J Hum Genet 32, 998–1004. 10.1038/s41431-024-01637-4.

17. Pagnamenta, A.T., Yu, J., Walker, S., Noble, A.J., Lord, J., Dutta, P., Hashim, M., Camps, C., Green, H., Devaiah, S., et al. (2024). The impact of inversions across 33,924 families with rare disease from a national genome sequencing project. Am J Hum Genet 111, 1140–1164. 10.1016/j.ajhg.2024.04.018.

18. Enver, T., Soneji, S., Joshi, C., Brown, J., Iborra, F., Orntoft, T., Thykjaer, T., Maltby, E., Smith, K., Abu Dawud, R., et al. (2005). Cellular differentiation hierarchies in normal and culture-adapted human embryonic stem cells. Hum Mol Genet 14, 3129–3140. 10.1093/hmg/ddi345.

19. Thompson, O., von Meyenn, F., Hewitt, Z., Alexander, J., Wood, A., Weightman, R., Gregory, S., Krueger, F., Andrews, S., Barbaric, I., et al. (2020). Low rates of mutation in clinical grade human pluripotent stem cells under different culture conditions. Nat Commun 11, 1528. 10.1038/s41467-020-15271-3.

20. Yang, S., Cho, Y., and Jang, J. (2021). Single cell heterogeneity in human pluripotent stem cells. BMB Rep 54, 505–515. 10.5483/BMBRep.2021.54.10.094.

21. Halliwell, J., Barbaric, I., and Andrews, P.W. (2020). Acquired genetic changes in human pluripotent stem cells: origins and consequences. Nature reviews. Molecular cell biology 21. 10.1038/s41580-020-00292-z.

22. DuBose, C.O., Daum, J.R., Sansam, C.L., and Gorbsky, G.J. (2022). Dynamic Features of Chromosomal Instability during Culture of Induced Pluripotent Stem Cells. Genes 13, 1157. 10.3390/genes13071157.

23. Bishop, D.C., Clancy, L.E., Simms, R., Burgess, J., Mathew, G., Moezzi, L., Street, J.A., Sutrave, G., Atkins, E., McGuire, H.M., et al. (2021). Development of CAR T-cell lymphoma in 2 of 10 patients effectively treated with piggyBac-modified CD19 CAR T cells. Blood 138, 1504–1509. 10.1182/blood.2021010813.

24. Ozdemirli, M. (2024). Indolent CD4+ CAR T-Cell Lymphoma after Cilta-cel CAR T-Cell Therapy - PubMed. https://pubmed.ncbi.nlm.nih.gov/38865661/.

25. Steffin, D.H.M., Muhsen, I.N., Hill, L.C., Ramos, C.A., Ahmed, N., Hegde, M., Wang, T., Wu, M., Gottschalk, S., Whittle, S.B., et al. (2022). Long-term follow-up for the development of subsequent malignancies in patients treated with genetically modified IECs. Blood 140, 16–24. 10.1182/blood.2022015728.

26. Tsuchida, C.A., Brandes, N., Bueno, R., Trinidad, M., Mazumder, T., Yu, B., Hwang, B., Chang, C., Liu, J., Sun, Y., et al. (2023). Mitigation of chromosome loss in clinical CRISPR-Cas9-engineered T cells. Cell 186, 4567–4582.e20. 10.1016/j.cell.2023.08.041.

27. Zou, L., Chen, Q., Quanbeck, Z., Bechtold, J.E., and Kaufman, D.S. (2016). Angiogenic activity mediates bone repair from human pluripotent stem cell-derived osteogenic cells. Sci Rep 6, 22868. 10.1038/srep22868.

28. Zhu, H., Blum, R.H., Bernareggi, D., Ask, E.H., Wu, Z., Hoel, H.J., Meng, Z., Wu, C., Guan, K.-L., Malmberg, K.-J., et al. (2020). Metabolic Reprograming via Deletion of CISH in Human iPSC-Derived NK Cells Promotes In Vivo Persistence and Enhances Anti-tumor Activity. Cell Stem Cell 27, 224–237.e6. 10.1016/j.stem.2020.05.008.

29. Cousineau, I., and Belmaaza, A. (2011). EMSY overexpression disrupts the BRCA2/RAD51 pathway in the DNA-damage response: implications for chromosomal instability/recombination syndromes as checkpoint diseases. Mol Genet Genomics 285, 325–340. 10.1007/s00438-011-0612-5.

30. Hughes-Davies, L., Huntsman, D., Ruas, M., Fuks, F., Bye, J., Chin, S.-F., Milner, J., Brown, L.A., Hsu, F., Gilks, B., et al. (2003). EMSY links the BRCA2 pathway to sporadic breast and ovarian cancer. Cell 115, 523–535. 10.1016/s0092-8674(03)00930-9.

31. Brakta, S., Hawkins, Z.A., Sahajpal, N., Seman, N., Kira, D., Chorich, L.P., Kim, H.- G., Xu, H., Phillips, J.A., Kolhe, R., et al. (2023). Rare structural variants, aneuploidies, and mosaicism in individuals with Mullerian aplasia detected by optical genome mapping. Hum Genet 142, 483–494. 10.1007/s00439-023-02522-8.

32. Costa, S.S., Fishman, V., Pinheiro, M., Rodrigueiro, A., Sanseverino, M.T., Zielinsky, P., Carvalho, C.M.B., Rosenberg, C., and Krepischi, A.C.V. (2024). A germline chimeric KANK1-DMRT1 transcript derived from a complex structural variant is associated with a congenital heart defect segregating across five generations. Chromosome Res 32, 6. 10.1007/s10577-024-09750-2.

33. Finlay, D., Murad, R., Hong, K., Lee, J., Pang, A.W.C., Lai, C.-Y., Clifford, B., Burian, C., Mason, J., Hastie, A.R., et al. (2024). Detection of Genomic Structural Variations Associated with Drug Sensitivity and Resistance in Acute Leukemia. Cancers (Basel) 16, 418. 10.3390/cancers16020418.

34. Neveling, K., Mantere, T., Vermeulen, S., Oorsprong, M., van Beek, R., Kater-Baats, E., Pauper, M., van der Zande, G., Smeets, D., Weghuis, D.O., et al. (2021). Next-generation cytogenetics: Comprehensive assessment of 52 hematological malignancy genomes by optical genome mapping. Am J Hum Genet 108, 1423–1435. 10.1016/j.ajhg.2021.06.001.

35. Sahajpal, N.S., Barseghyan, H., Kolhe, R., Hastie, A., and Chaubey, A. (2021). Optical Genome Mapping as a Next-Generation Cytogenomic Tool for Detection of Structural and Copy Number Variations for Prenatal Genomic Analyses. Genes (Basel) 12, 398. 10.3390/genes12030398.

36. Bosch, A., and Chillon, M. (2020). Gene therapy approaches in central nervous system regenerative medicine. In Handbook of Innovations in Central Nervous System Regenerative Medicine (Elsevier), pp. 375–398. 10.1016/B978-0-12-818084-6.00010-6.

37. Carter, M., and Shieh, J. (2015). Chapter 11 - Gene Delivery Strategies. In Guide to Research Techniques in Neuroscience (Second Edition), M. Carter and J. Shieh, eds. (Academic Press), pp. 239–252. 10.1016/B978-0-12-800511-8.00011-3.

38. Kalidasan, V., Ng, W.H., Ishola, O.A., Ravichantar, N., Tan, J.J., and Das, K.T. (2021). A guide in lentiviral vector production for hard-to-transfect cells, using cardiac-derived c-kit expressing cells as a model system. Sci Rep 11, 19265. 10.1038/s41598-021-98657-7.

39. Dixon-Salazar, T., Silhavy, J.L., Marsh, S.E., Louie, C.M., Scott, L.C., Gururaj, A., Al-Gazali, L., Al-Tawari, A.A., Kayserili, H., Sztriha, L., et al. (2004). Mutations in the AHI1 gene, encoding jouberin, cause Joubert syndrome with cortical polymicrogyria. Am J Hum Genet 75, 979–987. 10.1086/425985.

40. Yusa, K. (2015). piggyBac Transposon. Microbiol Spectr 3, MDNA3-0028-2014. 10.1128/microbiolspec.MDNA3-0028-2014.

41. Jinek, M., Chylinski, K., Fonfara, I., Hauer, M., Doudna, J.A., and Charpentier, E. (2012). A programmable dual-RNA-guided DNA endonuclease in adaptive bacterial immunity. Science 337, 816–821. 10.1126/science.1225829.

42. Komor, A.C., Kim, Y.B., Packer, M.S., Zuris, J.A., and Liu, D.R. (2016). Programmable editing of a target base in genomic DNA without double-stranded DNA cleavage. Nature 533, 420–424. 10.1038/nature17946.

43. Anzalone, A.V., Randolph, P.B., Davis, J.R., Sousa, A.A., Koblan, L.W., Levy, J.M., Chen, P.J., Wilson, C., Newby, G.A., Raguram, A., et al. (2019). Search-and-replace genome editing without double-strand breaks or donor DNA. Nature 576, 149–157. 10.1038/s41586-019-1711-4.

44. Miller, D.T., Adam, M.P., Aradhya, S., Biesecker, L.G., Brothman, A.R., Carter, N.P., Church, D.M., Crolla, J.A., Eichler, E.E., Epstein, C.J., et al. (2010). Consensus Statement: Chromosomal Microarray Is a First-Tier Clinical Diagnostic Test for Individuals with Developmental Disabilities or Congenital Anomalies. Am J Hum Genet 86, 749–764. 10.1016/j.ajhg.2010.04.006.

45. Langlois, S., Duncan, A., SOGC GENETICS COMMITTEE, and CCMG PRENATAL DIAGNOSIS COMMITTEE (2011). Use of a DNA method, QF-PCR, in the prenatal diagnosis of fetal aneuploidies. J Obstet Gynaecol Can 33, 955–960. 10.1016/S1701-2163(16)35022-8.

46. Shinawi, M., and Cheung, S.W. (2008). The array CGH and its clinical applications. Drug Discov Today 13, 760–770. 10.1016/j.drudis.2008.06.007.

47. Mardis, E.R. (2017). DNA sequencing technologies: 2006-2016. Nat Protoc 12, 213–218. 10.1038/nprot.2016.182.

48. Hor, P., Goldenson, Benjamin, and Kaufman, D.S. (2022). iPSC-Derived Natural Killer Cell Therapies - Expansion and Targeting. Front Immunol 13, 841107. 10.3389/fimmu.2022.841107.

49. Knorr, D.A., Ni, Z., Hermanson, D., Hexum, M.K., Bendzick, L., Cooper, L.J.N., Lee, D.A., and Kaufman, D.S. (2013). Clinical-Scale Derivation of Natural Killer Cells From Human Pluripotent Stem Cells for Cancer Therapy. Stem Cells Translational Medicine 2, 274–283. 10.5966/sctm.2012-0084.

50. Zhu, H., and Kaufman, D.S. (2019). An Improved Method to Produce Clinical-Scale Natural Killer Cells from Human Pluripotent Stem Cells. In In Vitro Differentiation of T-Cells: Methods and Protocols, S. Kaneko, ed. (Springer), pp. 107–119. 10.1007/978-1-4939-9728-2_12.

51. Kong, Y., Fierro, M., Pouyanfard, S., and Kaufman, D.S. (2022). Targeted Genomic Insertion of Cars in iPSC-Derived Macrophages Leads to Improved Expression and Anti-Tumor Activity. Blood 140, 7390–7391. 10.1182/blood-2022-159752.

52. Mardiros, A., Dos Santos, C., McDonald, T., Brown, C.E., Wang, X., Budde, L.E., Hoffman, L., Aguilar, B., Chang, W.-C., Bretzlaff, W., et al. (2013). T cells expressing CD123-specific chimeric antigen receptors exhibit specific cytolytic effector functions and antitumor effects against human acute myeloid leukemia. Blood 122, 3138–3148. 10.1182/blood-2012-12-474056.

53. Gill, S., Tasian, S.K., Ruella, M., Shestova, O., Li, Y., Porter, D.L., Carroll, M., Danet-Desnoyers, G., Scholler, J., Grupp, S.A., et al. (2014). Preclinical targeting of human acute myeloid leukemia and myeloablation using chimeric antigen receptor-modified T cells. Blood 123, 2343–2354. 10.1182/blood-2013-09-529537.

54. Li, X.-H., Goldenson, B., Thangaraj, J.L., Gynn, M., Gumber, D., Do, M., Willert, K., and Kaufman, D.S. (2022). Abstract 559: Targeting hematological malignancies and solid tumors with switchable chimeric antigen receptor-engineered iPSC-derived natural killer cells. Cancer Research 82, 559. 10.1158/1538-7445.AM2022-559.

55. Li, Y., Hermanson, D.L., Moriarity, B.S., and Kaufman, D.S. (2018). Human iPSC-Derived Natural Killer Cells Engineered with Chimeric Antigen Receptors Enhance Anti-tumor Activity. Cell Stem Cell 23, 181–192.e5. 10.1016/j.stem.2018.06.002.

56. Wang, J., Chen, S., Xiao, W., Li, W., Wang, L., Yang, S., Wang, W., Xu, L., Liao, S., Liu, W., et al. (2018). CAR-T cells targeting CLL-1 as an approach to treat acute myeloid leukemia. Journal of Hematology & Oncology 11, 7. 10.1186/s13045-017-0553-5.

